# Intraspecific genetic variation for anesthesia success in a New Zealand freshwater snail

**DOI:** 10.1101/2020.07.09.194050

**Authors:** Qiudong Song, Richard Magnuson, Joseph Jalinsky, Marissa Roseman, Maurine Neiman

## Abstract

Intraspecific genetic variation can drive phenotypic variation even across very closely related individuals. Here, we demonstrate that genetic differences between snails are a major contributor to wide variation in menthol anesthesia success in an important freshwater snail model system, *Potamopyrgus antipodarum*. Anesthesia is used to immobilize organisms for experiments and surgical procedures and to humanely mitigate pain. This is the first example of which we are aware of a role for genetic variation in anesthesia success in a mollusk. These findings highlight the fact that using only one strain or lineage for many experiments will not provide a full picture of phenotypic variation, demonstrate the importance of optimizing biomedically relevant techniques and protocols across a variety of genetic backgrounds, illuminate a potential mechanism underlying previously documented challenges in molluscan anesthesia, and set the stage for powerful and humane manipulative experiments in *P. antipodarum*.

## INTRODUCTION

While the establishment of genotype-phenotype links remains one of the outstanding open questions in biology (Mackay and Huang 2018; Baier et al. 2019; National Academies of Sciences 2020), there is clear consensus that genetic variation drives important behavioral (e.g., Niepoth and Bendesky 2020), neurological (e.g., Trimmer et al. 2019), physiological (e.g., Faralli and Lawson 2020; Gibney 2020), and life history (e.g., Takou et al. 2019) traits. Here, we demonstrate the existence of genetic variation for the phenotype of anesthesia efficacy in *Potamopyrgus antipodarum*, a New Zealand freshwater snail.

Anesthesia efficacy is a trait of interest to many both for its ethical and biomedical importance and as well as its existence at the nexus of physiology and neurobiology. From an ethical perspective, although animal welfare regulations often exclude invertebrates at least in part because how invertebrates process pain is not well understood (Cooper 2011; Gilbertson and Wyatt 2016; AMVA 2020), invertebrates nevertheless exhibit clear responses to stimuli that could be perceived as perception of pain (AVMA 2020). Indeed, Canada, the United Kingdom, and the European Union have animal research regulations that extend protections to cephalopods (Butler-Struben et al. 2018), potentially signifying a paradigm shift regarding which taxa deserve formal protections. The basis for animal welfare regulation of cephalopod research is the assumption, rooted in observational studies (e.g., Wells 1978; Roper and Hochberg 1988), that cephalopods perceive pain and exhibit emotional responses. This assumption has since been backed up empirically (e.g., Scientific Panel on Animal Health and Welfare 2005; Crook et al. 2013).

From the perspective of the biomedical community, anesthesia efficacy is important because researcher perception of the wellbeing of a research organism can affect study outcomes. For example, as outlined by Poole (1997), conclusions reached from studies using organisms under distress may be unreliable, meaning that pain experienced by research organisms can influence study results and conclusions (also see AVMA 2020).

Other than cephalopods, there are no other general protections extended to invertebrates for use in research of which we are aware. In the absence of regulatory oversite, given the likelihood that at least some invertebrates perceive pain, and because some experimental procedures are facilitated by or require that the animal is immobilized, there is a clear demand for the ability to reliably and effectively apply anesthesia in invertebrates.

We here describe important steps forward in identifying an effective anesthesia technique for *Potamopyrgus antipodarum*, a New Zealand freshwater snail model system that is a prominent model system for the evolution of sexual reproduction (e.g., Neiman et al. 2011), ecotoxicology (e.g., Geiß et al. 2017), invasion biology (e.g., Donne et al. 2020), and host-parasite interactions (e.g., Bankers et al. 2017). Studies of *P. antipodarum* often incorporate tissue manipulation (e.g., tentacle severing as a test of tissue regeneration; Krois et al. 2013) or require determination of male vs. female status (e.g., Jalinsky et al. 2020). These studies either require or would be qualitatively improved by the availability of an effective, safe, and easy-to-use anesthetic.

Menthol crystals are the preferred method for anesthetizing *P. antipodarum* (e.g., Krois et al. 2013), with a standard approach based on exposure of snails to crushed menthol crystals for ∼90 minutes (McCraw 1958). While this method can be effective (Krois et al. 2013), a considerable fraction (often >50%) of exposed *P. antipodarum* do not become anesthetized (Magnuson 2018). Similar intraspecific variation and generally low anesthesia efficiency has been reported in other mollusks (e.g., Fiorito et al. 2015; Butler-Struben et al. 2018).

Magnuson (2018) noted that some *P. antipodarum* lineages (defined as all snails descended from a single, laboratory-isolated female) seemed to have markedly higher anesthesia success than others. This observation led us to hypothesize that this lineage effect points to an important role of genetic background in anesthesia success in these snails. This hypothesis finds indirect support from reports of genetically based differences in anesthesia efficacy in rodents (e.g., Chesler et al. 2003; Moghil et al. 2005; Seltzer 2014), insects (e.g., Guan et al. 2000), and roundworms (Morgan and Sedensky 1994; Hawasli et al. 2004; reviewed in Nash 2002; Steele et al. 2007).

Here, we performed the first test of which we are aware of a genetic basis for anesthesia efficacy in mollusks by subjecting different genetic lineages of common garden-born and raised *P. antipodarum* asexual lineages to the same menthol anesthesia protocol. The presence of significance across-lineage variation in anesthesia success would be consistent with this hypothesis, suggesting that some *P. antipodarum* genetic backgrounds are more susceptible to menthol anesthesia than others. This result will be useful to other *P. antipodarum* and mollusk researchers going forward, highlighting a potential explanation for frequently reported challenges in anesthesia success and indicating that at least some experiments requiring anesthesia can succeed if susceptible lineages are used. More broadly, our findings highlight the importance of including a diversity of strains or lineages when optimizing methods or developing protocols that interface with phenotypes that plausibly feature intraspecific genetic variation.

## METHODS

### Anesthesia experiment

We followed the definition of successful anesthesia provided by Lewbart and Mosley (2011), who considered the snail to be anesthetized when the animal shows no tentacle withdrawal or body movement following the gentle scrape of its foot with a needle (“needle test”). For the experiments presented herein, we applied the anesthetization protocol used for *P. antipodarum* by Krois et al. (2013). The process begins with the addition of 200 mL of room-temperature carbon-filtered tap water to a ∼1/2-L plastic cup held in a room-temperature laboratory. We then added the *P. antipodarum* to be anesthetized to the cup, immediately followed by the addition of 2 g of crushed menthol crystals. After 90 minutes of exposure, we removed the snails from the menthol water. We defined anesthesia as successful if the body of the snail was relaxed such that the tentacles, head, and foot had emerged from the shell and there was no withdrawal or body movement during the needle test (administered with a 25-gauge needle). We deemed anesthesia as unsuccessful if the snail was fully contracted into the shell or if the snail was relaxed but failed the needle test.

We then administered this anesthesia protocol to each of six groups of ten snails (N = 60 snails per lineage) from each of 17 different *P. antipodarum* triploid asexual lineages, for a total of 102 trials and 1020 snails (Table 1). While each of the 17 lineages was founded by a single unique asexual female and was cultured separately in its own tank, three pairs of lineages (six of the 17 lineages) shared descent from the same field-collected grandmother. These founding grandmothers were sampled from natural populations in 2017, about two generations before the birth of the snails used here. Snails, lineages, and lakes were chosen as a combined function of snail availability and the existence in culture of multiple lineages per lake. While we cannot formally rule out the possibility that transgenerational effects of lake of origin (apart from genetic variation *per se*) are still influencing snail phenotypes, we believe that it is reasonable to expect that several generations of culture in a standardized laboratory environment should provide ample time to minimize such effects.

**Table 1.** Description of the *p. antipodarum* asexual lineages used in this study.

| Lake of Origin | Lineage | Grandmother | Mean Anesthesia Success % (+/- SD) | Genotyped | SNP Genotype |
| --- | --- | --- | --- | --- | --- |
| Alexandrina | Alex 13 | Alex 13 | 23.33 +/- 15.01 | N | NA |
| Alexandrina | Alex 19_1 | Alex 19 | 50.00 +/- 6.32 | Y | 3 |
| Alexandrina | Alex 19_2 | Alex 19 | 53.33 +/- 10.33 | Y | 3 |
| Alexandrina | Alex 21_1 | Alex 21 | 15.00 +/- 8.37 | Y | 2 |
| Alexandrina | Alex 21_2 | Alex 21 | 35.00 +/- 12.25 | N | NA |
| Alexandrina | Alex 27 | Alex 27 | 91.67 +/- 11.69 | Y | 4 |
| Alexandrina | Alex 31 | Alex 31 | 48.33 +/- 7.53 | N | NA |
| Alexandrina | Alex 32 | Alex 32 | 95.00 +/- 8.37 | Y | 4 |
| Alexandrina | Alex 36 | Alex 36 | 95.00 +/- 5.48 | Y | 5 |
| Alexandrina | Alex 39 | Alex 39 | 93.33 +/- 8.16 | Y | 4 |
| Grasmere | Grasmere 49_1 | Grasmere 49 | 76.67 +/- 15.01 | Y | 26 |
| Grasmere | Grasmere 49_2 | Grasmere 49 | 48.33 +/- 9.83 | Y | 26 |
| Grasmere | Grasmere 54 | Grasmere 54 | 83.33 +/- 13.66 | Y | 26 |
| Grasmere | Grasmere 59 | Grasmere 59 | 8.33 +/- 7.53 | Y | 24 |
| Gunn | Gunn 6 | Gunn 6 | 36.67 +/- 20.66 | Y | 29 |
| Gunn | Gunn 7 | Gunn 7 | 71.67 +/- 16.02 | Y | 26 |
| Gunn | Gunn 9 | Gunn 9 | 78.33 +/- 14.72 | Y | 31 |

We expected that these three pairs of lineages (each pair defined by descent from the same grandmother) would be genetically identical (barring *de novo* mutations) across pair members (hereafter, “clonemates”). We were able to back up these assumptions by leveraging single-nucleotide polymorphism (SNP) data (see “SNP genotyping”). By enabling comparisons between clonemates (N = three pairs) as well as across genetically distinct lineages (“non-clonemates”; 14 lineages, including those three lineage pairs), our set of lineages thus allows us to test whether genetic factors influence anesthesia success.

All of the snails in our experiment were born and raised in the same constant-temperature room, under identical feeding and culture conditions (following standard of care for *P. antipodarum*; e.g., Zachar and Neiman 2013). The snails used for the trials were haphazardly selected from the adult snails in the 15 L tanks used to house each lineage and were not reused across trials. Our measure of anesthesia success per trial was the percentage of snails, out of the ten snails per lineage per trial, that were successfully anesthetized.

Because the anesthesia success data were not distributed normally (Kolmogorov-Smirnov test; test statistic = 0.111, df = 102, *p* = 0.003), we used a non-parametric Kruskal-Wallis test to first address whether there was a significant effect of lineage founder (“grandmother”) on anesthesia success. A significant outcome of this Kruskal-Wallis analysis will indicate that the descendants of different grandmothers respond differently to anesthesia. Because the founding grandmothers constituted separately sampled snails from highly diverse New Zealand populations (e.g., Paczesniak et al. 2013; Verhaegen et al. 2018), this outcome is consistent with a major role of genetic variation for anesthesia success in *P. antipodarum*.

We next used the lineage-level anesthesia success data to address whether the three pairs of clonemates were significantly more similar in anesthesia success rates than were non-clonemate pairs. We began by calculating the absolute value of the difference in anesthesia success rates (“difference scores”) between the means of the six success rates measured for each of the 17 lineages relative to each of the other lineages. For example, this value for Alex 13 relative to Alex 27 is the absolute value of 0.23 (Alex 13 mean success rate) - 0.92 (Alex 27 mean success rate), equivalent to 0.69. There were a total of three unique pairwise difference scores for the clonemates (reflecting the three pairs of clonemates) and a total of 136 unique pairwise difference scores for the non-clonemate comparisons. We then used one-sample sign tests (as implemented at https://www.mathcelebrity.com/nonparam.php?pl=Generate+Practice+Problem) relative to a test value of zero, to determine whether the clonemates and non-clonemate difference scores, respectively, were significantly different than zero. In a scenario where genetic background is an important determinant of anesthesia success, the clonemate difference score would not be significantly different than zero, while the non-clonemate score should be significantly higher than zero. Except where indicated otherwise, all statistical analyses and figures were executed and drawn, respectively, in IBM SPSS Statistics v. 25.

### SNP Genotyping

As part of a larger and distinct population genomics project, we used KASP™ assays to genotype 14 of the 17 lineages at 49 of the 50 SNPs used in Verhaegen et al. (2018) (Online Resource 1). We began by using the same chloroform-phenol DNA extraction method described in Sharbrough et al. (2018) on the dissected head tissue of one randomly selected adult snail from each of the fourteen lineages. We then sent the DNA extractions to LGC Genomics (Hoddesdon, UK) for SNP genotyping. Upon receiving the raw data, we assigned SNP genotypes by using the infinite alleles model distance index, setting the maximum distance threshold defining a genotype (Rogstad et al. 2002) to zero differences (excluding the three sites of 686 total genotyped for which the genotype was not able to be called) with GenoDive software v.2.0 (Meirmans and Van Tienderen 2004).

## RESULTS

Our analyses revealed wide across-lineage variation in anesthesia outcome between lineages descended from different grandmothers (Kruskal-Wallis, *p* < 0.001; Figure 1), ranging from a median of 95% of all snails anesthetized for several different lineages from lake Alexandrina to ∼8% for Grasmere 59 (Table 1). This result is as is expected if there is a major genetic component to anesthesia success in *P. antipodarum*.

**Fig. 1.**
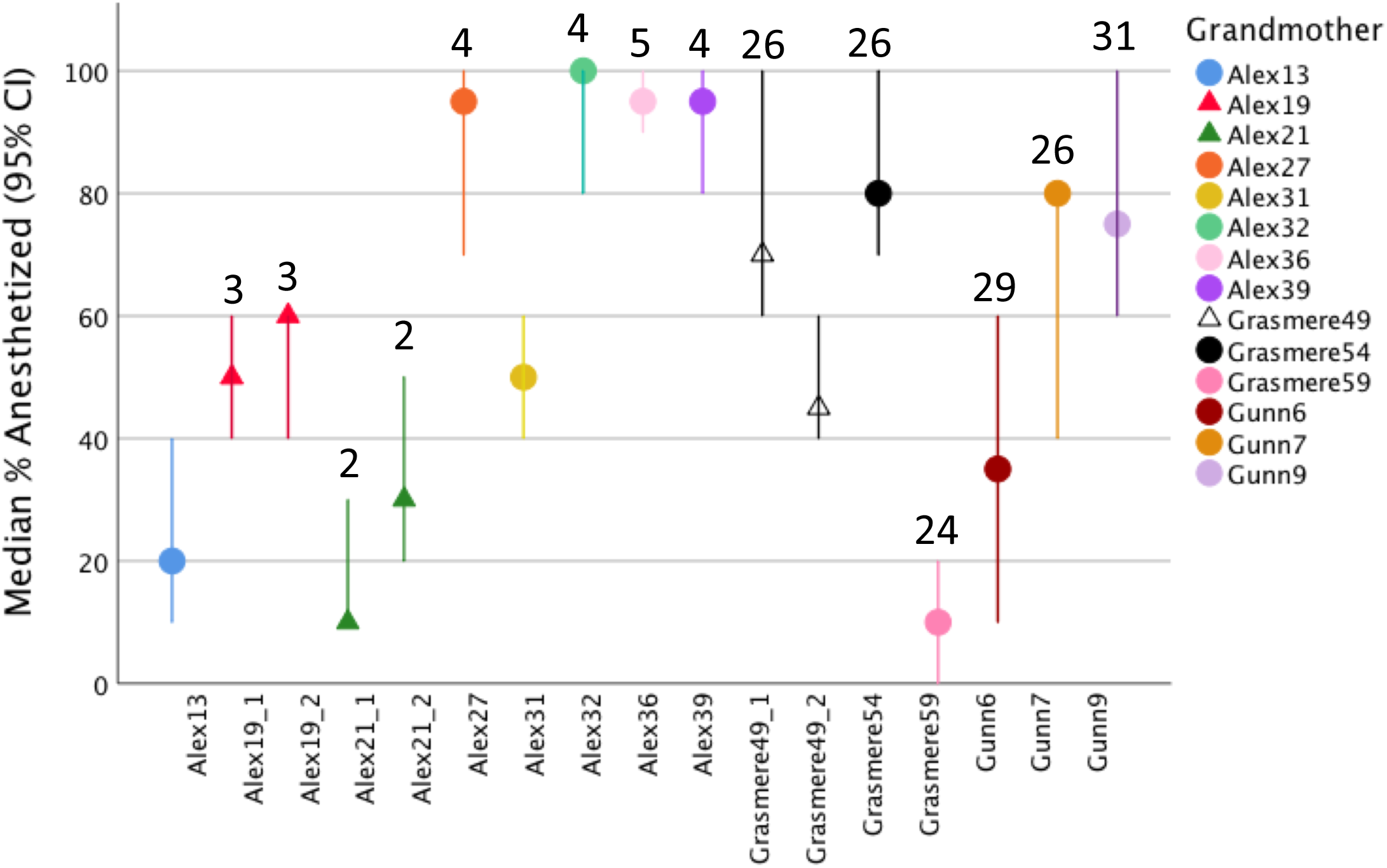
Median percent of successfully anesthetized snails (6 trials of 10 snails each) subjected to the standard *P. antipodarum* menthol anesthesia protocol per each of 17 lineages founded by 14 different grandmothers. Triangle symbols (vs. circles) are used to denote the three lineage pairs that each share a grandmother. The numbers above the 14 of the 17 lineages that were genotyped indicate SNP genotype, with shared numbers indicating shared SNP genotypes

The difference score between clonemates was not significantly different than zero (one-sample sign test; *p* = 0.125), with the caveat that we had only three comparisons and thus low statistical power. By contrast, the difference score between non-clonemates was significantly higher than zero (*p* = 0.002) and was nearly twice as high (mean difference score 0.340 +/- 0231 SD) as the mean difference score for the clonemates (mean difference score = 0.173 +/- 0.132 SD; Fig. 2). These results are also consistent with a scenario where genetic factors influence anesthesia success in *P. antipodarum*, though a firm conclusion awaits validation from a follow-on experiment with substantially more statistical power.

**Fig. 2.**
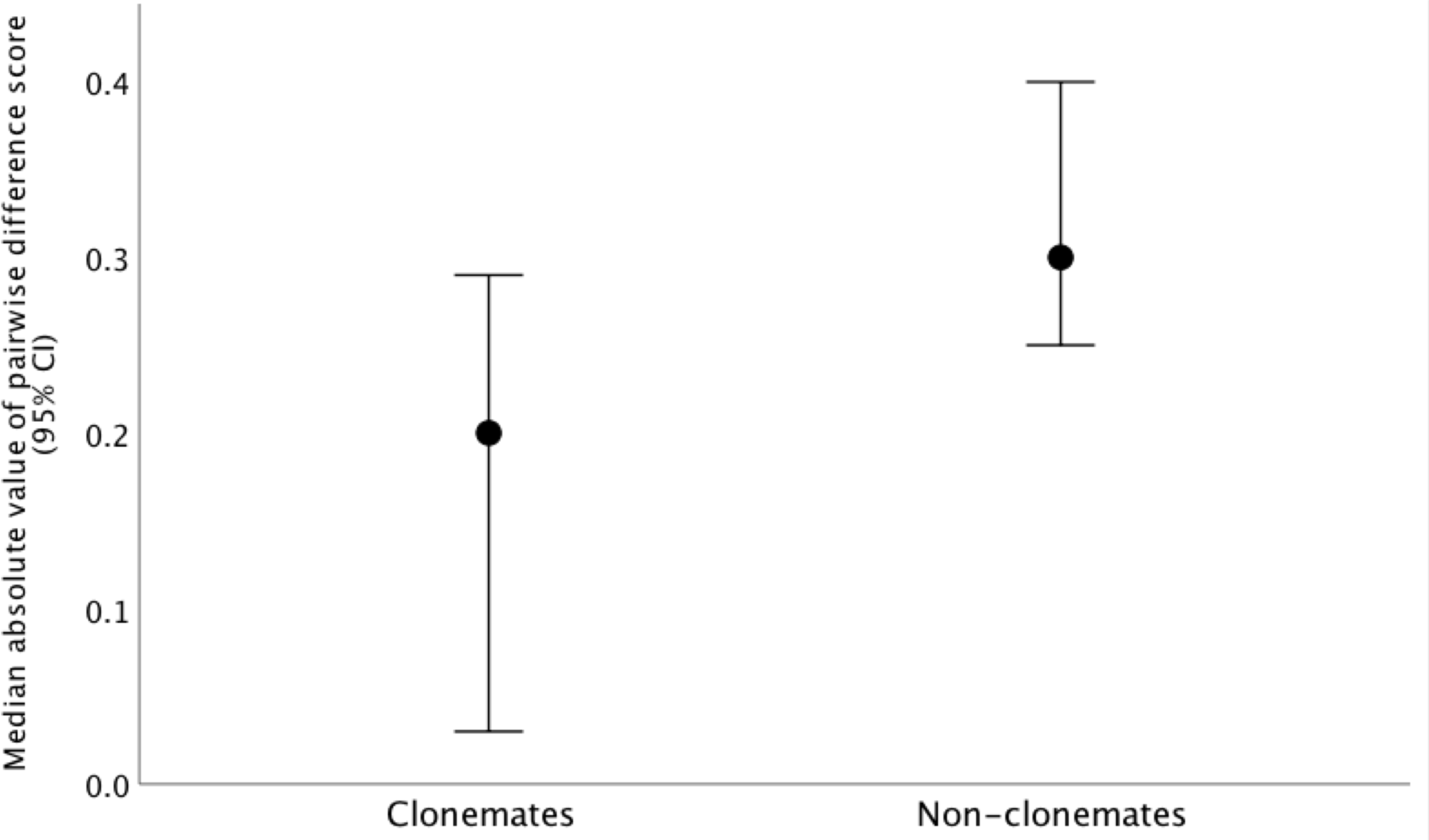
Median pairwise difference score in anesthesia success for clonemate vs. non-clonemate lineages. The non-clonemate pairwise difference scores were significantly higher than zero while the clonemate pairwise difference scores were not significantly higher than zero

There were eight distinct SNP genotypes across the 14 lineages genotyped (Table 1). These SNP data demonstrated that the two pairs of clonemates that were genotyped (Alex 19_1 and Alex 19_2 and Grasmere 49_1 and Grasmere 49_2) indeed shared SNP genotypes. These data also revealed that most other non-clonemate lineages themselves had unique genotypes. Finally, we discovered that the two sets of lineages with different grandmothers that nevertheless shared SNP genotypes 4 and 26, respectively, had strikingly similar anesthesia success rates relative to other lineages with different SNP genotypes (Fig. 1).

## DISCUSSION

Together, our data indicate that genetic background is a major contributor to menthol anesthesia success in *P. antipodarum*, an important model system in evolutionary biology, ecology, and ecotoxicology. This is the first demonstration of which we are aware of a genetic component to variation in anesthesia efficacy in mollusks. One obvious practical implication of these results for *P. antipodarum* researchers is the opportunity to select lineages with higher anesthesia susceptibility for experiments that require anesthetics. That genetic background influences anesthesia success in a wide variety of common model systems like flies, worms, and mice suggests that the same might be expected for other mollusk taxa. Considered even more broadly, our findings emphasize the critical importance of including representative intraspecific genetic diversity when optimizing methods or protocols for a model organism.

We were able to perform a powerful independent validation of the likelihood that genetic variation does underlie across-lineage variation in anesthesia success in *P. antipodarum* by leveraging single-nucleotide polymorphism (SNP) data generated for a different population genomic project. These data backed up our assumptions about the genetic identity of clonemates relative to non-clonemates and also revealed that lineages not previously known to be close relatives that shared a SNP genotype seemed to have very similar anesthesia outcomes relative to lineages with different SNP genotypes, though the strength of this conclusion is limited by low statistical power.

Two important questions worthy of future study emerge from our results. First, what is the genetic basis of the across-lineage variation in anesthesia success? Second, even though clonemates seem to share similar anesthesia success outcomes relative to non-clonemates, why do we not see identical responses (i.e., no examples of 100% success or 100% failure) within clonemates?

The first question can be addressed by using techniques such as RNA sequencing and whole-genome sequencing to identify gene expression differences and genetic variants differentiating lineages with very distinct anesthesia responses. Because multiple studies indicate that asexual *P. antipodarum* lineages from the same lake population are typically closely related relative to asexual lineages from other lakes (e.g., Paczesniak et al. 2013), a powerful experiment might be one that compared genomic sequence from lineages from the same lake with different anesthesia responses. A good example of such lineages in the experiment we describe here is provided by low-responding lineages from lake Alexandrina like Alex 13 (23.33% anesthesia success) and Alex 21 (15% anesthesia success) as compared to Alex 27 (91.67% anesthesia success) and Alex 32 and 36 (95% anesthesia success).

The second question, regarding variation in anesthesia success even within clones and clonemates, hints that other factors (e.g., size, age, body condition) could influence anesthesia outcomes (e.g., Larsson and Wahlström 1998; Guenette and Lair 2006; Gao et al. 2018). While we made a distinct effort to minimize age and size effects by choosing the largest snails possible from each lineage, we did not know the ages of individual snails. Nor are we able to exclude the possibility that across-snail differences in size, health, or body condition could have come into play.

We were able to provide an initial assessment of a potential role for snail size/age by quantifying the length (measuring the longest possible distance between apex and aperture under a dissecting microscope) of each of 30 haphazardly selected individuals from each of the 60 individuals used for anesthesia trials for each lineage. This value is meaningful for *P. antipodarum* because the snails increase in body length until reproductive maturity, though across-lineage variation in adult size means that some adult snails are larger than others (Larkin et al. 2016). We then used a Spearman’s correlation analysis to determine whether the mean body length per lineage was associated with the median anesthesia success for each lineage.

We predicted that a strong effect of age or size on our results would manifest in a relationship between body length and anesthesia success outcomes. Not only did this analysis not reveal any such relationship (Spearman’s correlation coefficient = -0.06, *p* = 0.82; Fig. 3), but the two Alex 19 lineages, with mean body lengths differing by over a millimeter (likely reflecting different demographic structure in the source tanks) nevertheless had virtually identical anesthesia success. This result, though preliminary, does provide an independent line of support that genetic differences across lineages are a main driver of differences in anesthesia success rates. Future experiments could address a role for size, body condition, or age more directly by comparing anesthesia success within a lineage as a function of snail age or in the context of experimental manipulation of factors like food availability.

**Fig 3.**
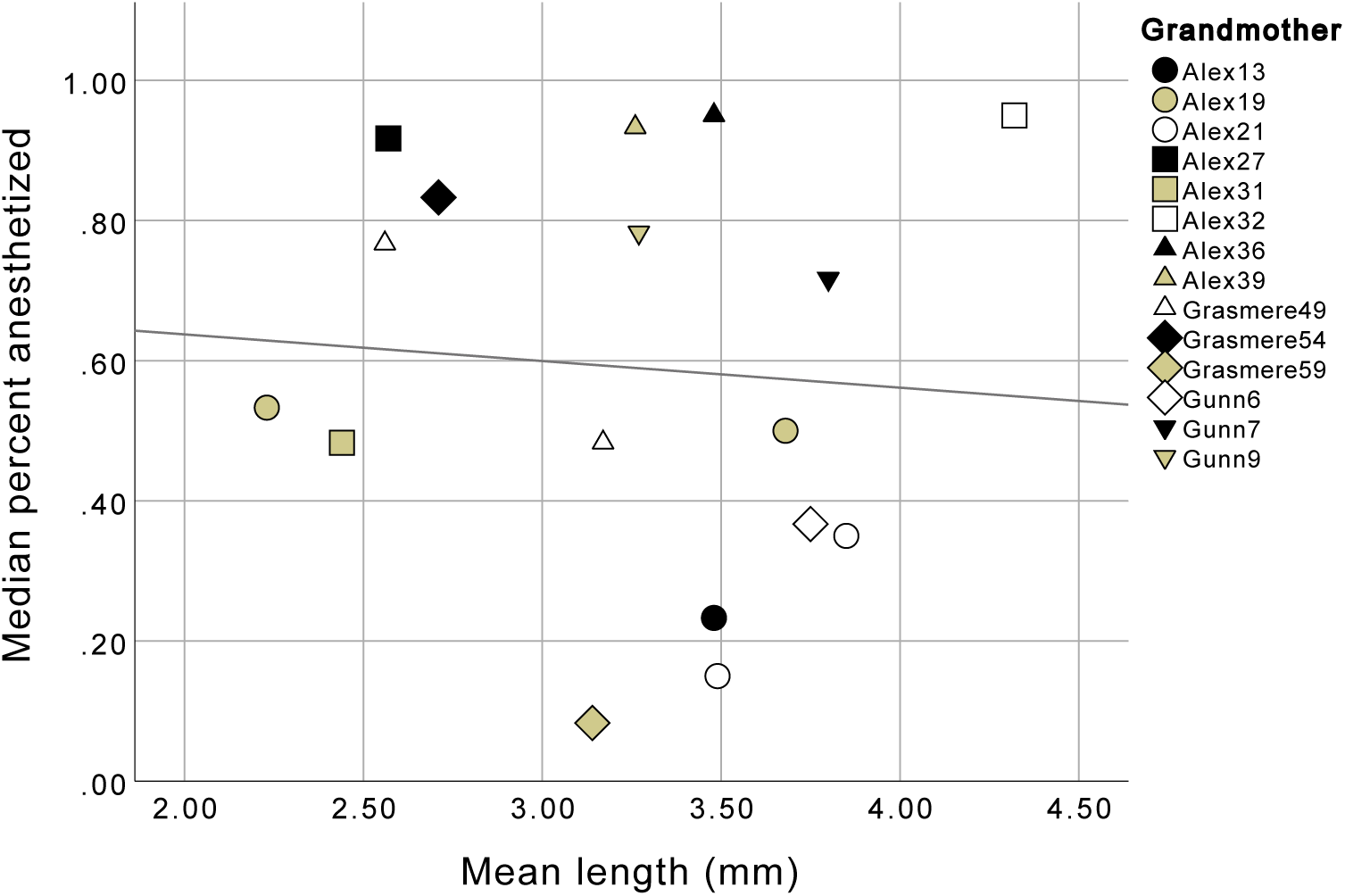
Mean size per lineage (mm) vs. median anesthesia success rate (percent) for each of the 17 lineages. The black diagonal line represents the best-fit linear relationship between the two variables

## Supporting information

Electronic supplemental material 1

## Acknowledgements

We gratefully acknowledge Molly Gallagher, Sydney Stork, Benjamin Ripperger, and Alexander Kern for snail care. John Logsdon contributed to snail collections. David (“Davey”) Neiman contributed his Excel expertise to data analysis. Several anonymous reviewers of an earlier version of the manuscript provided valuable feedback. The University of Iowa Department of Biology Honors program, the Iowa Center for Undergraduate Research, and Rick and Linda Maxson logistically and financially supported several undergraduate researchers involved in the project. The SNP genotyping described here was also supported by National Science Foundation - MCB 1122176, National Science Foundation - DEB 1731657, and National Science Foundation - DEB 1601242.

## Declarations

### Ethics approval

Not applicable.

### Consent to participate

Not applicable.

### Consent for publication

Not applicable.

### Availability of data and material

Data will be made available upon request.

### Code availability

Not applicable

### Authors’ contributions

Maurine Neiman, Richard Magnuson, and Qiudong Song designed the study. Richard Magnuson and Qiudong Song collected the snail phenotype data. Joseph Jalinsky and Marissa Roseman generated the SNP data. Maurine Neiman analyzed the phenotype data and Joseph Jalinsky analyzed the SNP data. Maurine Neiman, Richard Magnuson, and Qiudong Song wrote the manuscript, and Joseph Jalinsky and Marissa Roseman provided editorial revisions. All authors approved the final manuscript.

## Electronic Supporting Material

**ESM_1** SNP markers genotyped for each of 14 lineages. “?” indicates that the marker for that snail could not be called accurately

